# Reversible peptide self-assembly enables sustained drug delivery with tuneable pharmacokinetics

**DOI:** 10.64898/2026.03.25.714189

**Authors:** Therese W. Herling, Jiapeng Wei, Sivaneswary Genapathy, Cristian Rivera, Marie Persson, Peter Gennemark, David Workman, Dan Lundberg, Elise Bernard, Hannah Bolt, Marianna Yanez Arteta, Sarah Will, Annette Bak, David Hornigold, Tuomas P. J. Knowles, Ana L. Gomes dos Santos

## Abstract

Therapeutic peptides combine high target specificity with potent biological activity.^1^ However, treatment success is often limited by rapid clearance and the need for frequent injections.^2, 3^ This challenge is particularly acute for therapeutic peptides used in obesity, where clinical benefit must be balanced against dose-dependent adverse effects. In nature, these constraints are overcome by storing hormones as reversible fibrils,^4^ but pharmacokinetic control is essential for widespread adoption of bio-inspired self-assembled depots for therapeutic peptides. Here, we show that tuneable pharmacokinetics can be achieved and modelled by mapping the fundamental chemical parameters of reversibly self-assembly *in vitro*. We demonstrate this approach for the amylin analogue pramlintide. Amylin analogues are under development for the next generation of diabetes and obesity treatments, with improved mechanism of action e.g. preserving lean body mass.^5–8^ Pramlintide is an approved drug with a well-established safety profile, however, it has a comparable half-life to native amylin.^8–12^ In a pilot study, we achieve *in vitro*–*in vivo* correlation, increasing the half-life of pramlintide 20–82-fold in rats, while controlling burst release. These findings demonstrate that the optimisation of pharmacokinetics can be decoupled from peptide engineering, establishing a generalisable framework for generating long-acting peptide formulations by emulating native storage mechanisms.

---

Peptide therapeutics have driven a paradigm shift in the treatment of chronic metabolic conditions and play a pivotal role in improving patient outcomes. For over a century, insulin has transformed type 1 diabetes from a fatal condition to a manageable one.^13^ Worldwide, the need for effective and accessible peptide-based treatments is growing; over one billion people live with obesity and 830 million people have diabetes.^14, 15^ Incretins such as glucagon-like peptide-1 (GLP-1) and amylin analogues collectively address the underlying pathophysiology of these metabolic diseases.^5, 16^ However, native-like peptides are prone to chemical and enzymatic degradation, in addition to rapid renal clearance due to their small size, Figure 1**a**.^2, 3^ The pharmacokinetics of therapeutic peptides can be extended by modifying the peptide itself e.g. reducing proteolytic lability or renal clearance with peptide conjugates.^17, 18^ However, the addition of a bulky conjugate can compromise receptor binding, reducing drug potency.^5, 19^ The optimisation of GLP-1 based peptide-lipid conjugates for weekly dosing has revolutionised the treatment of diabetes and obesity.^16, 20^ Long-acting formulations are therefore key to unlocking the full potential of peptide therapeutics.

**Figure 1:**
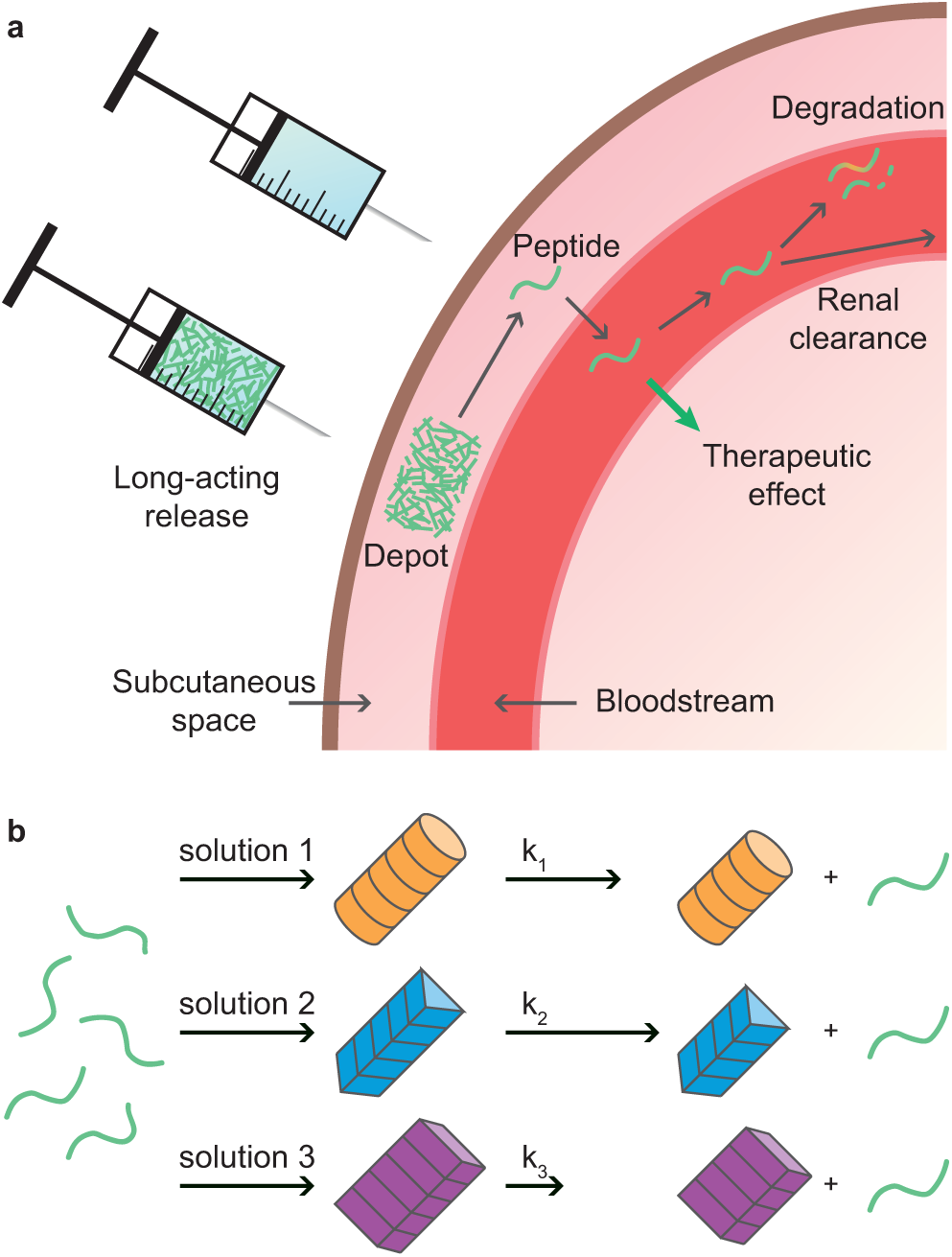
Sustained peptide delivery. **a** Unmodified peptides are prone to fast renal clearance, chemical and biochemical degradation, limiting their half-life *in vivo*, typically minutes to hours.^2, 3, 25^ Peptide self-assembly can be used for an all-peptide depot to deliver functional peptide over days or weeks.^10–12, 26^ **b** We use the solution environment for the self-assembly reaction to tune the morphology and release kinetics of peptide depots. This approach decouples the optimisation of pharmacokinetics from the peptide chemistry to generate long-acting peptide formulations without compromising on drug efficacy.

We focus on pramlintide in this study. The 37 amino acid peptide is an amylin receptor agonist and approved for the treatment of type 1 and type 2 diabetes.^21, 22^ Amylin analogues have the potential to improve on current weight management treatments by limiting the loss of lean mass, which can lead to frailty in patients following GLP-1 only treatments.^8^ However, pramlintide has a similar half-life to native amylin, requiring multiple daily injections.^9, 21–23^ Long-acting amylin analogues are under development, including as combination therapies with GLP-1.^5, 8, 24^ Strategies that could extend the pharmacokinetics of these peptides are therefore highly desirable in the current pharmaceutical landscape.

As an alternative to modifying the peptide itself, a polymer or lipid depot matrix can be employed to sterically trap, store and release therapeutic peptide.^17, 27, 28^ However, peptide release from depot matrices show complex kinetics e.g. poly-(D,L-lactide-co-glycolide) (PLGA).^29^ Initial burst release of loosely associated peptide can produce transient concentration spikes that limit the tolerable injectable dose. Poor predictive power for *in vivo* pharmacokinetics based on *in vitro* data means that optimisation of the drug-depot combination requires considerable resources.^30^

In biological function, peptides and proteins self-assemble to form fibrils with a cross-*β* structure typical of the amyloid fold. These assemblies have functions in signalling, regulation of cellular processes, storage and release, and as scaffolds.^31, 32^ Naturally occurring fibrils that act as storage depots for controlled release of active proteins or peptides, including peptide hormones,^4^ are of great interest in the context of long-acting peptide therapeutics.^26^ This approach is employed for the somatostatin analogue lanreotide, yielding dosing intervals of 4-8 weeks.^10–12^ Self-assembly into fibril structures has also been demonstrated for the drugs degarelix and oxyntomodulin, extending the lifetime of oxyntomodulin in rats from four hours to five days.^26, 33^ Indeed, most proteins and peptides can self-assemble given the right solution conditions, and the fibrillar form is one of the most thermodynamically stable protein conformations,^34, 35^ Bio-inspired self-assembly is therefore a potential general-purpose strategy for generating long-acting peptide formulations.

Tuneable pharmacokinetics are needed to deliver a generalisable approach to the design of self-assembled formulations. A major challenge for this concept is that the dissociation environment is fixed, as peptide release occurs in the subcutaneous space. Here, we program the peptide depot pharmacokinetics by varying the self-assembly conditions, and we demonstrate that different self-assembled structures deliver distinct thermodynamics and kinetic profiles. We use *in vitro*-determined parameters to model *in vivo* serum concentrations and burst release. This framework decouples pharmacokinetic optimisation from peptide primary structure and medicinal chemistry, thereby preserving intrinsic receptor potency and selectivity while achieving tailored exposure profiles. This approach has the potential to remove major barriers to treatment compliance and uptake of peptide therapeutics.

## Results and Discussion

### Regulating peptide self-assembly through the buffer composition

First, we identify favourable self-assembly conditions for pramlintide by performing a screen across pH, buffer system, and ionic strength at 37^◦^C, Figure 2 and SI Figure 1. Therapeutic pramlintide acetate (Symlin) is formulated at a concentration of 0.6–1 mg/ml in acetate buffer at pH 4 with D-mannitol for isotonicity.^36^ We accelerate pramlintide self-assembly by adding NaCl instead of D-mannitol to increase the ionic strength, SI Figure 1. The appearance of fibril material is monitored through the reporter dye Thioflavin T (ThT).^37^ Fluorescence time courses were acquired for pramlintide concentration series with a maximum of 10 mg/ml (2.53 mM) with 150 mM NaCl in seven buffers (30 mM), Figure 2**a** and Methods. 0.01% v/v Tween 20 was added to the samples used for kinetic analysis to prevent peptide adsorption at interfaces. In Figure 2**a-b**, we include lower pramlintide concentrations for pH 6-7.4, see *m_t_* range in panel **b**. We found that pramlintide self-assembled across the pH range investigated here (pH 3-7.4), and that the reaction accelerates with increasing pH.

**Figure 2:**
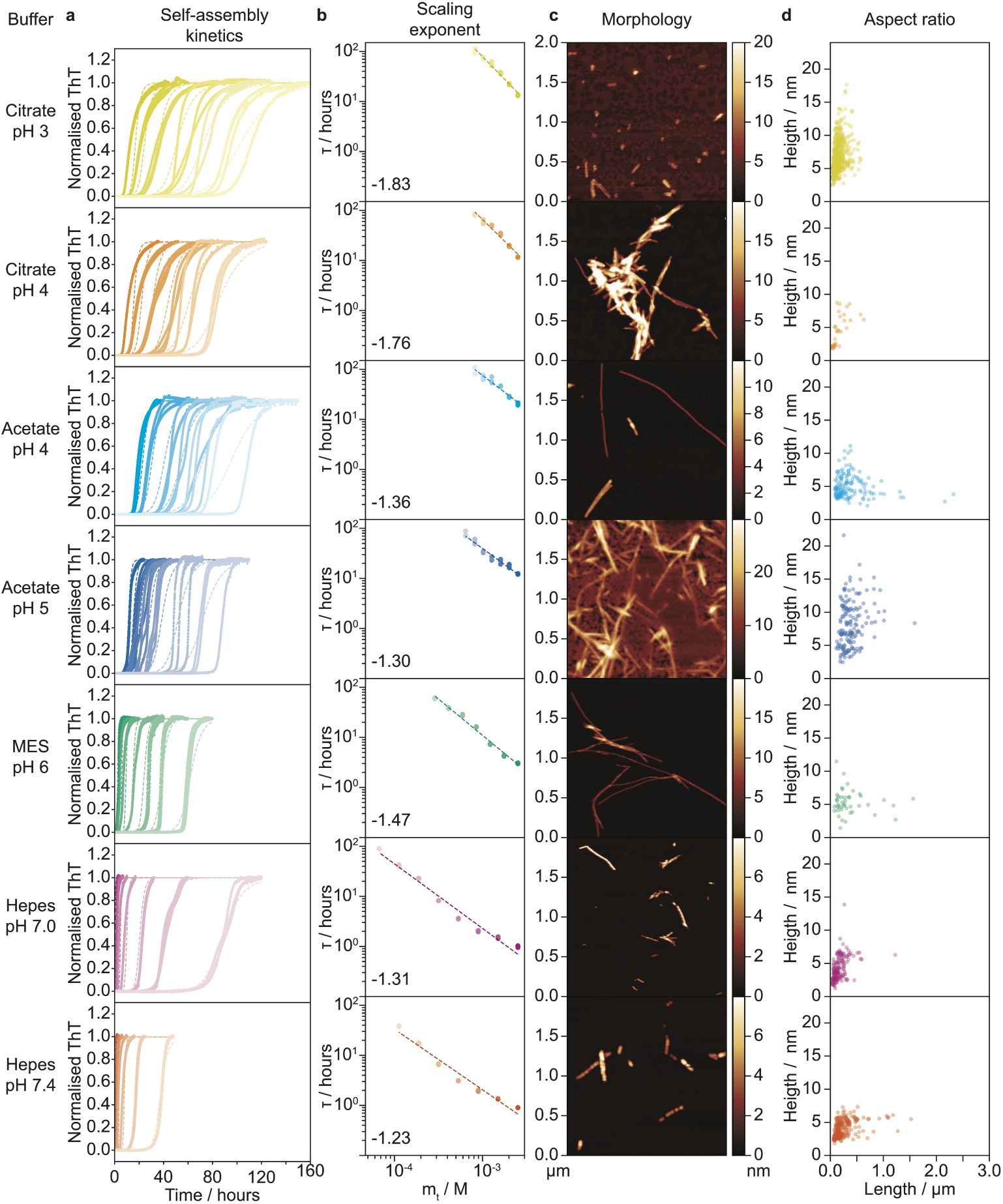
The pramlintide self-assembly reaction accelerates as a function of pH. **a** Pramlintide self-assembly monitored by ThT and normalised to the plateau fluorescence. Global fits (dashed lines) for each buffer were made using a model with primary nucleation, elongation and secondary nucleation, setting *n_c_* = 2.^37^. **b** Scaling exponents and half-time (*τ*) versus total peptide concentration. The range of concentrations assayed was adapted to the overall reaction speed for each buffer. **c** AFM images of the fibril structures show distinct morphologies. **d** The fibril dimensions depend on the self-assembly buffer. The aspect ratio can be used to tune the number of fibril ends per unit peptide monomer-equivalent in the reaction.

To understand the underlying molecular mechanisms, we examined the scaling exponent (*γ*), Figure 2**b**, which provides information on the dominating processes in the self-assembly reaction.^37, 38^ For pramlintide, the scaling exponents were consistent with a secondary nucleation model (*γ* = -(*n*_2_+1)/2), where the reaction is accelerated by secondary nucleation events catalysed by existing fibrils, and a secondary nucleus size *n*_2_ ≈2, Figure 2**b**.^37, 38^ We performed global fits to the kinetic data for each buffer condition as detailed in Methods with three free global parameters: *k*_+_*k_n_*, *n*_2_, *k*_+_*k*_2_. Where *k*_+_ is the elongation rate, *k_n_* the primary nucleation rate, and *k*_2_ the secondary nucleation rate. The primary nucleus size, *n_c_*, was fixed as 2, Table 1 and SI Figure 2**a-b**. Here, we find that self-assembly accelerates with pH through secondary processes.

**Table 1:** Critical concentrations, average fibril lengths, and rate constants (*n_c_* = 2) for pramlintide as a function of the buffer system.

| Buffer | $c_c / \mu\text{M}$ | $l / \mu\text{m}$ | Scaling | $n_2$ | $k_+ / \text{M}^{-1} \text{hr}^{-1}$ | $k_- / \text{hr}^{-1}$ | $k_n / \text{M}^{-n_c} \text{hr}^{-1}$ | $k_2 / \text{M}^{-n_2} \text{hr}^{-1}$ |
| --- | --- | --- | --- | --- | --- | --- | --- | --- |
| Citrate pH 3 | $27.2 \pm 22.5$ | $0.15 \pm 0.09$ | -1.83 | 2.52 | $3.2 \cdot 10^2$ | $8.8 \cdot 10^{-3}$ | $9.7 \cdot 10^{-4}$ | $1.1 \cdot 10^6$ |
| Citrate pH 4 | $17.8 \pm 13.3$ | $0.36 \pm 0.16$ | -1.76 | 2.26 | $3.1 \cdot 10^4$ | $5.4 \cdot 10^{-1}$ | $8.3 \cdot 10^{-6}$ | $2.9 \cdot 10^3$ |
| Acetate pH 4 | $12.7 \pm 8.90$ | $0.39 \pm 0.35$ | -1.36 | 1.70 | $4.7 \cdot 10^4$ | $6.0 \cdot 10^{-1}$ | $6.2 \cdot 10^{-6}$ | $2.5 \cdot 10^1$ |
| Acetate pH 5 | $11.0 \pm 8.94$ | $0.40 \pm 0.21$ | -1.30 | 1.91 | $7.5 \cdot 10^4$ | $8.3 \cdot 10^{-1}$ | $3.0 \cdot 10^{-6}$ | $3.2 \cdot 10^2$ |
| MES pH 6 | $0.308 \pm 0.258$ | $0.38 \pm 0.33$ | -1.47 | 1.75 | $2.9 \cdot 10^4$ | $8.8 \cdot 10^{-3}$ | $1.2 \cdot 10^{-6}$ | $5.9 \cdot 10^3$ |
| Hepes pH 7 | $0.593 \pm 0.395$ | $0.18 \pm 0.14$ | -1.31 | 1.99 | $1.6 \cdot 10^5$ | $9.4 \cdot 10^{-2}$ | $2.8 \cdot 10^{-5}$ | $1.9 \cdot 10^5$ |
| Hepes pH 7.4 | $0.227 \pm 0.125$ | $0.26 \pm 0.22$ | -1.23 | 2.17 | $2.9 \cdot 10^5$ | $6.7 \cdot 10^{-2}$ | $2.7 \cdot 10^{-6}$ | $9.4 \cdot 10^5$ |

We investigated the nanoscale morphology of self-assembled pramlintide using atomic force microscopy (AFM), Figure 2**c**, and in the monomer hydrodynamic radius by microfluidic diffusional sizing, SI Figure 3. Short, wide fibrils are formed in citrate pH 3, whereas long, slim fibrils are formed in acetate pH 4, MES pH 6, and Hepes buffers. These differences in fibril height (*h*) and length (*l*), Figure 2**d**, result in variation in the number of fibrils (*f* = *M/l*) and growing ends (*n_end_* = 2*f*) per unit monomer concentration, Table 1. We integrate *n_end_* in our analysis of the individual rate constants.

**Figure 3:**
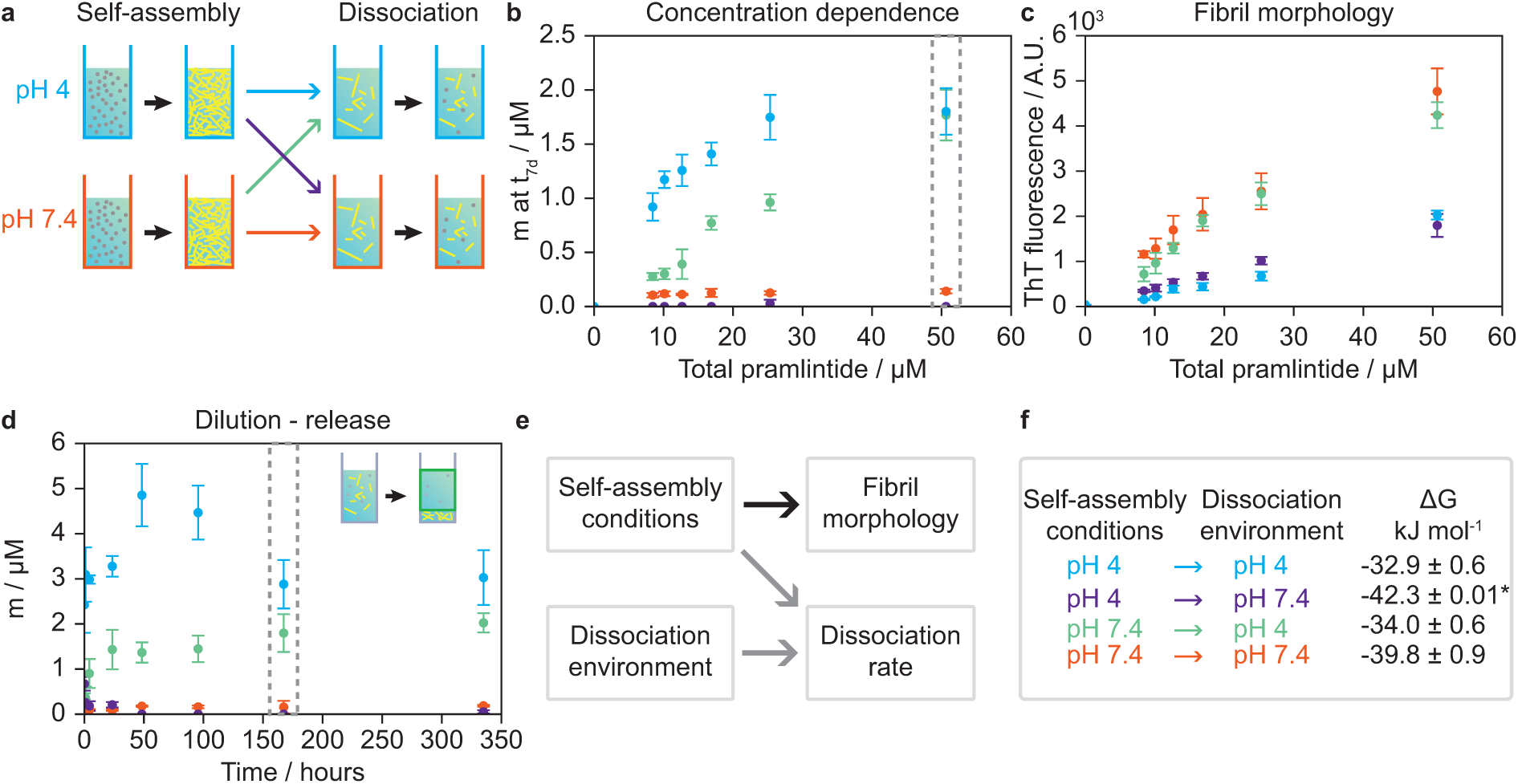
Tuneable dissociation kinetics for pramlintide. **a** Schematic of experiment workflow, 10 mg/ml pramlintide is self-assembled in acetate pH 4 or Hepes pH 7.4, both with 150 mM NaCl, 20 *µ*M ThT, 0.01% v/v Tween20. The two fibril types are then diluted in either pH 4 or pH 7.4 buffer to create four sample sets, two with the dilution buffer is different from the self-assembly buffer, and two where the solution conditions match. The legend in the bottom right corner of the figure applies to all panels. **b** Soluble pramlintide as a function of *m_t_* at *t* = 7 days. Grey dashed boxes show equivalent data points in **b,d**. **c** ThT fluorescence intensity at t = 7 days as a function of pramlintide concentration. The ThT fluorescence intensity depends on the fibril structure. **d** Soluble pramlintide as a function of time following dilution of 10 mg/ml fibril solutions to 0.2 mg/ml (51 *µ*M) in acetate pH 4. **e** The self-assembly conditions dominate fibril morphology, whereas both self-assembly and dissociation environment regulate peptide release. **g** Legend and ΔG for the four buffer combinations based on *m* = *c_c_* for 7-14 days (panel **d**). ^∗^ Only two samples with *m* above the detection limit in panel **d**.

In addition to measuring the formation of fibril material, we also determined the endpoint concentration of free monomer by ultra performance liquid chromatography (UPLC), Table 1 and SI Figure 2**c**. The endpoint concentration of soluble peptide reflects the critical concentration (*c_c_*) for a given peptide and solution environment.^35^ At equilibrium,

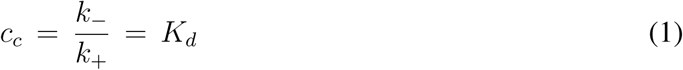

we were therefore able to gain information on the equilibrium parameters. For pH 3-5, the endpoint free monomer concentration is 11-27 *µ*M, whereas for pH 6-7.4 it is only 0.2-0.6 *µ*M. The concentration of soluble peptide is important for depot formulations, where monomer release soon after injection can lead to burst kinetics.^29^ A peak in plasma peptide concentration (high *c_max_* with short *t_max_*) may surpass the therapeutic window, increasing the risk of adverse effects and reducing tolerability. Unlike matrix-based depots, the soluble monomer concentration (*m*) has a constant value when the total pramlintide concentration *m_t_* ≫ *c_c_*. As an example, for a 10 mg/ml solution, the free peptide would be 0.009±0.005% of the total dose for Hepes pH 7.4, which could be reduced further by increasing *m_t_*. Optimising *c_c_* during formulation development can limit burst release of the therapeutic peptide to generate a favourable pharmacokinetic profile.

Traditionally, peptide and protein fibrils are viewed as persistent structures, with negligible dissociation rates,^37, 38^ limited research has therefore focussed on reversible self-assembly.^39^ Nonetheless, fibril dissociation is an important process for hormone release,^4^ disease-related aggregate clearance,^40^ and the release of therapeutic peptide from self-assembled depots.^12, 26, 41^ Due to the practical challenges of quantifying the relatively small amount of peptide that is released over long timescales, the errors on experimentally determined *k*_−_ values typically exceed those for measurements of *k*_+_. We therefore approach *k*_−_ through the forward reaction by determining *k*_+_ from the growth of preformed fibrils and *K_d_*, Equation 1, Table 1 and SI Figures 2&4.^42^ From *k*_+_ and *K_d_*, we determined *k*_−_ values for the seven buffer systems in this study. We decided to focus on acetate pH 4, where pramlintide is chemically stable to e.g. de-amidation,^9, 10^ and pH 7.4 in the physiological range. Comparing these two conditions, pramlintide self-assembly is more favourable at pH 7.4 with a lower *c_c_*and *k*_−_, as well as faster fibril elongation, *k*_+_.

### Differences in self-assembly conditions translate to tuneable dissociation kinetics

We have shown that the self-assembly environment leads to fibrils with distinct morphologies, reaction kinetics and equilibrium parameters, Table 1. Next, we investigate whether peptide release depends on the fibril characteristics, the dissociation environment, or a combination of these factors. To test our hypothesis, that we can use the self-assembly conditions to encode the peptide release kinetics, we generate a cross of self-assembly and dissociation conditions for acetate pH 4 and Hepes pH 7.4, Figure 3**a**.

Here, we investigated peptide release as a function of *m_t_* after dilution (10 mg/ml stocks to 0.03-0.2 mg/ml, 8.4-51 *µ*M), Figure 3**b-c** and SI Figure **5**. Both the fibrillar, *M*, and soluble peptide, *m*, are diluted at time (*t*) = 0 days, leading to a soluble pramlintide concentration, *m* ≠ *c_c_*. The sample then re-equilibrates to reach *c_c_*for the dilution conditions. We collected the soluble pramlintide fraction at *t* = 7 days and measured the peptide concentration by UPLC, Figure 3**b** and Methods. For pH 7.4→7.4, the concentration of soluble peptide is constant against *m_t_*, here 0.12 ± 0.03 *µ*M, which is consistent with *c_c_*from the forward kinetics, Table 1 and SI Figure 2. The soluble peptide concentration for pH 4→7.4 is near or below the detection limit (0.06 *µ*M), with one sample at 0.08 *µ*M. Changing the solution conditions for pH 4 type fibrils therefore lowers *c_c_* from 12.7 *µ*M to ≤0.08 *µ*M.

The ThT fluorescence yield varies between fibril isoforms, with a higher fluorescence intensity for pH 7.4 fibrils, Figure 2**c**. After seven days, the ThT fluorescence intensity of the fibrils still group by their formation conditions, Figure 3**c**, indicating that the fibril morphology is conserved, even when the solution environment is changed, a feature, we also observe by AFM, SI Figure 6.

Unlike pH 7.4→7.4 fibrils, the endpoint concentrations of soluble peptide for fibrils diluted in pH 4 buffer increase with *m_t_*. This correlation could be due to incomplete re-equilibration at *t* = 7 days or low polymerisation cooperativity, as observed for cytoskeletal protein filaments.^43, 44^ To investigate the time scales of re-equilibration following dilution, we monitored peptide release from 1 in 50 dilutions to (0.2 mg/ml, 51 *µ*M), Figure 3**d**. Here, we find that plateau values are reached by *t* = 2 days, with most of the peptide release/uptake occurring within the first hours. In both Figure 3**b,d**, we measure *c_c_* ≈2-3 *µ*M for fibrils at pH 4, which is lower than the endpoint monomer concentration measured for the forward reaction, Table 1. This observation is not unusual, as dense fibril networks can limit monomer access to growing ends and fibril surfaces for secondary nucleation, meaning that *m* reaches *c_c_* on a longer timescale than the plateau in ThT fluorescence. We therefore verify *c_c_* for self-assembly conditions of interest via dilution-dissociation experiments.^45^ We confirmed the *c_c_* values in Figure 3 by replacing the supernatants collected for analysis at each time point with blank buffer solution, the soluble fraction was then collected again at *t* = 14 days, SI Figure 5. When we compare data for peptide release as a function of *m_t_*and time, we observe lower release for pH 7.4→4 compared to pH 4→4 and fast uptake to pH 4→7.4 fibrils. Critically, both the self-assembly and dissociation environment contribute to the dissociation reaction.

We use *c_c_*for *t* = 7-14 days (Equation 1) to determine the change in Gibb’s free energy (Δ*G* = *R T ln*(1*/K_d_*)), and we find that for pH 4→7.4 fibrils are more thermodynamically stable than pH 7.4→7.4 fibrils, Figure 3**g**. We investigated, whether these differences in ΔG originated in changes to *k*_+_, *k*_−_, or both in a cross-seeding assay, SI Figure 7. Growth was slow for pH 7.4→4 fibrils, even compared to pH 4→4 fibrils. On the other hand, fibrils formed at pH 4 showed rapid growth when added to 1 mg/ml pramlintide, Hepes pH 7.4, with *k*_+_ = 3.3·10^5^ M^−1^ hr^−1^, corresponding to *k*_−_ = 0.025 hr^−1^. This comparison shows that the self-assembly environment can lead to distinct growth and release properties.

### The peptide self-assembly environment encodes pharmacokinetics *in vivo*

Having demonstrated that the self-assembly conditions for pramlintide can be used to tune peptide dissociation *in vitro*, Figure 3, we investigated whether this tuneable release translated to differential pharmacokinetics *in vivo*. Fibril dissociation depends on *n_end_* and *k*_−_, while elongation depends on *k*_+_, *n_end_*, and the concentration of free monomer (*m_t_*− *M* (*t*)).^38^ For the depot, we replace *m_t_* − *M* (*t*) with the subcutaneous peptide concentration *m_sc_*:

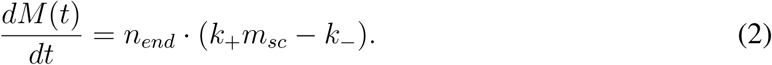

Depot release is combined with a two-compartment pharmacokinetic model (*V*_1_, *V*_2_) with elimination rate (*k_e_*), using a literature value for the absorption rate (*k_a_*), Figure 4**a**.^9, 46, 47^ Below, we report rates, concentrations and volumes for a rat with bodyweight = 0.3 kg.

**Figure 4:**
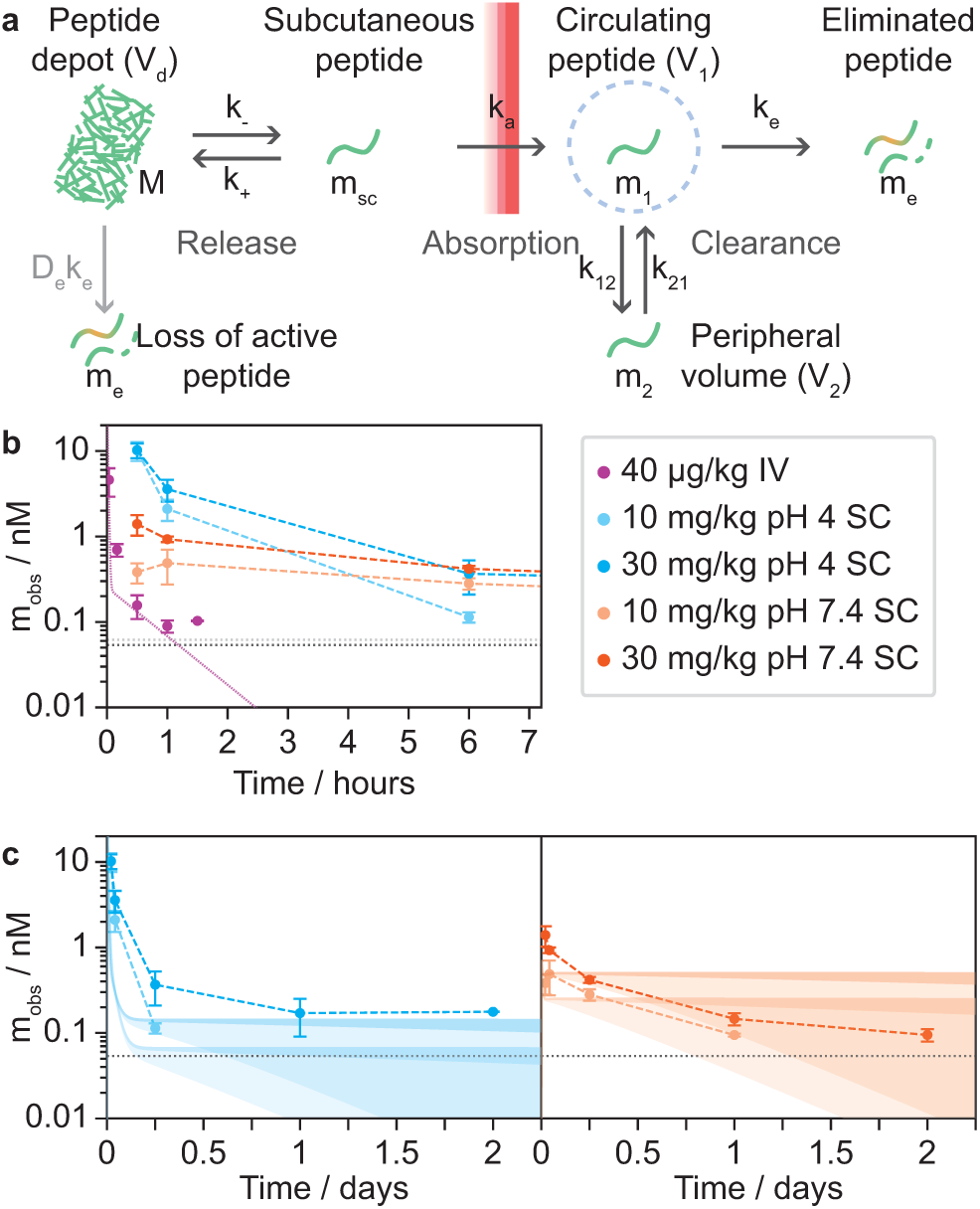
Tunable pharmacokinetics of release from subcutaneous depot. **a** Schematic of pramlintide release from depot, absorption, and clearance. **b** Serum pramlintide concentration (*m_obs_*) in rats following either a single injection of a subcutaneous depot or an intravenous infusion of pramlintide (see legend). The early times show fast clearance of IV peptide, and burst release of free monomer in pH 4 depot. A two-compartment model is fitted to the IV data, dotted purple line. Error bars show the standard error of the mean. At *t* = 1.5 h in the IV arm, 2 of 3 samples were ¡ LLOQ and were included in the model fit as LLOQ/2. Dotted horizontal lines the lower limit of quantification for the IV (light grey) and subcutaneous (dark grey) samples. Depot data points are connected by dashed lines. **c** Detectable pramlintide is released from the depot formulations over the course of days for the pH 4 depot (left) and pH 7.4 depot (right). Shaded areas are predicted peptide concentration in the central volume using the model in **a**, with *k*_+_ and *k*_−_ as predicted from the *in vitro* self-assembly, *k_e_*, *k*_12_, *k*_21_, *V*_1_, *V*_2_ from the IV data, and a literature value for *k_a_* = 8 hr^−1^,^46^ with the shaded areas covering *D_e_* values of 0 to 10^−4^ (dark) and 10^−3^ (light).

For the pharmacokinetic study, we chose two self-assembly conditions: acetate pH 4 and Hepes pH 7.4, Figure 4. The self-assembled formulations are relatively simple compared to PLGA or LiquidCrystal depots, consisting of peptide, NaCl and buffer.^30, 48^ The aim of this study was to characterise the pharmacokinetics of peptide release from a subcutaneous depot. Accordingly, we used lean (Sprague Dawley) rats rather than diet-induced obese animals, as the study was not designed to assess efficacy.

Four groups of rats received a single injection of a subcutaneous depot at doses of either 10 or 30 mg/kg self-assembled pramlintide, with two control groups receiving vehicle only (n = 4 per group), Figure 4**b**, Table 2 and Methods. Separately, serum pramlintide concentrations were recorded following IV administration of 40 *µ*g/kg non-assembled pramlintide in acetate pH 4.8 with D-mannitol for isotonicity. This IV bolus enters *V*_1_ directly as circulating peptide and enables us to fit *k*_12_ = 11.65 hr^−1^, *k*_21_ = 1.79 hr^−1^, *k_e_* = 92.5 hr^−1^ and the two compartment volumes *V*_1_ = 0.012 L, *V*_2_ = 0.078 L. The depot formulation concentrations were 5 and 15 mg/ml to give dose volume *V_d_*= 0.6 ml, Figure 4**a**. The self-assembled pramlintide depot was administered through a narrow-gauge needle (27G in this exploratory study) compared to e.g. LiquidCrystal (22G) and long-acting lanreotide (18G), minimising discomfort.^48^ In this study, the subcutaneous depots were well-tolerated, and we did not observe any inflammation or irritation at the injection site.

**Table 2:**
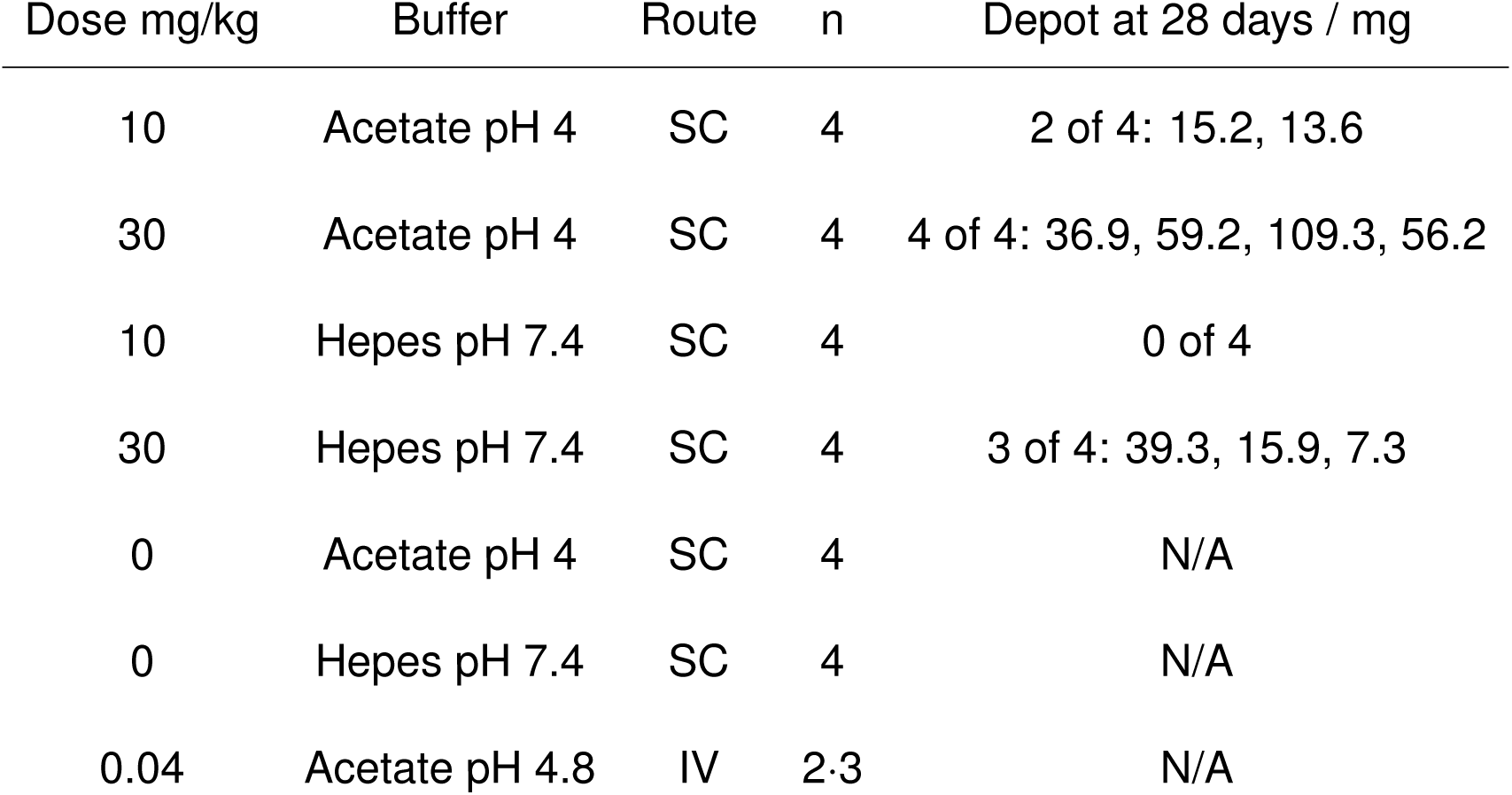
Benchmark pharmacokinetic study in lean rats. Four groups received a depot of self-assembled pramlintide through a single subcutaneous (SC) injection. Two groups received the vehicle only. Six rats received pramlintide intravenously (IV), these were divided into two groups of three for sample collection at alternating time points.

Our model predicts that *m_obs_*(Figure 4**a**) will initially be dominated by release of any soluble peptide in the depot, followed by a steady-state phase, and decrease with depot depletion. In the steady state phase, the local depot concentration *M* will be in the mM range, with *m_sc_*in the 10-100 nM range, and *m_obs_*= *m*_1_ in the 10-100 pM range, Figure 4**c**. Even with a relatively high *k_e_*= 92.5 hr^−1^, Figure 4**b**, peptide removal during the steady state phase occurs at ≈10^−8^ M hr^−1^. When accounting for the volume change between *V_d_*and *V*_1_, it would take ≈10^3^ hr to eliminate the depot via this pathway only. Peptide elimination from the subcutaneous space reduces bioavailability and has been included in previous models for therapeutic peptide depot formulations e.g. oxyntomodulin.^26^ We therefore consider a secondary elimination pathway from the depot environment, with the rate constant *D_e_*· *k_e_*, in our analysis below.

We found distinct pharmacokinetics for the two depot formulations, Figure 4**b**, demonstrating that the peptide pharmacokinetics can be tuned through the self-assembly conditions. A comparison of *t*_1*/*2_ between time points for the IV reference (*t*_1*/*2_ = 0.4 hr, consistent with literature values of *t*_1*/*2_ = 0.5-1 hr),^23, 46^ with the subcutaneous depots (*t*_1*/*2_ = 8.9-36 hr for *t* ≥ 6 hr) show a 20–82-fold increase in *t*_1*/*2_ for the depots. This result effectively decouples the development of the therapeutic peptide from the optimisation of the drug pharmacokinetics.

Based on the endpoint concentrations *in vitro*, Table 1, we would expect burst release for the depot formulations in acetate pH 4 (*c_endpoint_*= 12.7 ±8.9 *µ*M). Indeed, we observe an initial spike in serum pramlintide in rats, with a dose-independent *c_max_* = 10 nM, Figure 4**b-c**, followed by extended release from the depot. A 300 g rat would receive 7.6 nmol soluble peptide with the pH 4 formulation. When we average *m_obs_*for the first hour and combine with the clearance value of *CL* = 1.11 L hr^−1^ from the IV data, the initial peptide release for the pH 4 depots is 6.8 and 6.2 nmol for the 10 and 30 mg/kg doses respectively. These values are in excellent agreement with the predicted 7.6 nmol peptide,^9, 46^ considering that fibrils made at pH 4 will take up some monomer, when transferred to physiological pH, Figure 3. On the other hand, for the depots formulated in Hepes pH 7.4, where *c_c_* is 0.227 ± 0.125 *µ*M, burst release does not dominate the initial *m_obs_*, and *c_max_* shows dose-proportionality at 0.5 and 1.4 nM. This data demonstrates that we are able to predict not only which formulations will have burst release, but also the amount of peptide that is released during the burst phase.

We have shown that *k*_+_ and *k*_−_ for the reversible self-assembly reaction depend on both the self-assembly and dissociation environment *in vitro*, Table 1 and Figure 3. Next, we investigate the correlation between predictions based on our *in vitro* analysis and the observed pharmacokinetic data *in vivo* by performing a numerical integration of the ordinary differential equation for *m*_1_:

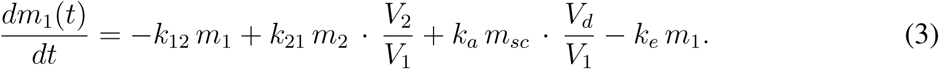

Here, we use *k*_+_ and *k*_−_ for pH 4→7.4 and pH 7.4→7.4, as the parameters for peptide release in the pH 7.4 buffer are closest to physiological conditions, SI Figure 8. Notably, we use parameters from our *in vitro* analysis of the self-assembly reaction to predict release from the depot formulations directly, rather than performing a fit of the model to the data, and we find a good agreement between the predicted and observed serum concentrations across different doses and formulation types. The model predicts 10^−8^ M concentrations for *m*_1_ (*m_obs_* in the study) at early times for the pH 4 formulations, followed by steady state concentrations of 10^−11^-10^−10^ M, while the pH 7.4 formulations are predicted in the 10^−10^ M range for both the early and steady state phases, Figure 4**c**. This *in vitro* - *in vivo* correlation enables the rational design of long-acting formulations, increasing the development that can be achieved through faster and less costly *in vitro* assays before progressing to *in vivo* testing.

In this analysis, we set *m_sc_*= *c_endpoint_*for *t* = 0 hr to account for the free peptide in the depot formulation, this approximation is likely to underestimate the duration of burst release following depot injection, as it does not account for the time it takes for the peptide to diffuse out of the depot matrix. Indeed, *m_obs_* is above steady state levels for the first hours of the study, which aligns with the duration of burst release from other depot types e.g. PLGA.^29^

The pharmacokinetic data shows a decrease in *m_obs_* over the course of days until it falls below the detection limit (54 pM). However, subcutaneous depot is still found in the majority of rats at *t* = 4 weeks, Table 2. These results indicate that peptide release continues below the lower limit of quantification (LLOQ) for weeks after a single injection, with the lower dose depots being depleted first. Without a secondary elimination pathway, the model predicts a near-constant *m_obs_* during the first days of the study, see upper bound of shaded area in Figure 4**c**. The measured slope in *m_obs_* suggests that the peptide depot is eliminated via a second pathway, and that this pathway reduces *M* directly, rather than eliminating *m_sc_*, which would decrease the overall *m_obs_*, as opposed to introducing a slope. We include depot elimination *D_e_* at a fraction of the elimination rate for circulating peptide. When we consider the amount of depot recovered at 28 days (7.3-109.3 mg from a ≈0.6 ml injection), we can constrain *D_e_*to the region 10^−4^, Figure 4**c** and SI Figure 9. This range for *D_e_* agrees with the *in vivo* data for the pH 7.4 formulations, where the depot environment has the highest similarity to the physiological conditions for peptide release. For the pH 4 formulations, the measured *m_obs_* is at the upper end of the predicted range, but then falls below the LLOQ at longer timescales (pramlintide detected in 2/4 serum samples at *t* = 2 days for 30 mg/kg). *D_e_* may vary between fibril structures and depot formulations, and this parameter could be fitted with additional data.

The well-controlled burst release from self-assembled depots enables high-dose depots. We simulate the serum pramlintide concentration as a function of dose, SI Figure 10. The results of these simulations highlight two design parameters for self-assembled depot formulations: the duration of peptide release and the steady state concentration of peptide as a function of dose. On one hand, *k*_−_ and *n_end_*are lower for the pH 4 depots, extending the release profile. On the other, the faster release from the pH 7.4 depots lead to higher *m_obs_* for a given dose. Self-assembly enables high depot doses to be delivered with comparable (pH 4) or lower (pH 7.4) *c_max_*to a much smaller IV bolus. For example, the predicted *c_max_* for a 100 mg/kg pH 7.4 depot is below the one observed for a 40 *µ*g/kg IV injection, Figure 4**b** and SI Figure 10**b,d**. In addition, for a 100 mg/kg depot, therapeutically relevant serum concentrations (≥ 21 pM)^49^ would be maintained for 2-3 weeks following a single injection, for *D_e_* = 10^−4^, SI Figure 10**c-d**.

Pramlintide is not a long-acting peptide, and is eliminated from circulation shortly after being released from the depot. However, if we consider our approach applied to modern long-acting amylin or GLP-1 analogues (e.g. cagrilintide *t*_1*/*2_ = 159-195 hr), and we vary *k_e_* to increase *t*_1*/*2_, our model predicts peptide release with *m*_1_ values near *c_max_*over several months, SI Figure 11. Self-assembled depots of e.g. long-acting lipidated peptides therefore have the potential for sustained delivery of therapeutic peptides on timescales that are significantly beyond the current state-of-the-art.^8^

### Conclusions

In this study, we address one of the main barriers to success for peptide therapeutics, the short half-lives of native-like peptides. We demonstrate that reversible self-assembly of an unmodified therapeutic peptide can be engineered to form bio-inspired depots with tuneable pharmacokinetics. We increase the half-life 20–82 fold in a formulation dependent manner. By analysing the underlying physicochemical parameters for the reversible self-assembly reaction, we show that both the self-assembly and dissociation environments regulate peptide release from depot structures. These findings establish the fundamental physicochemical principles for depots with distinct pharmacokinetics, but consisting of the same peptide. This work opens the door for rational design of long-acting peptide therapeutics with programmable pharmacokinetics by establishing *in vitro*-*in vivo* correlation for burst release and serum concentrations as a function of depot preparation. In this proof-of-concept study, we work with pramlintide, which has a half-life of 26 minutes, similar to native amylin, and we improve otherwise challenging pharmacokinetics. However, self-assembly is a common process accessible to most proteins and peptides. In combination with recently developed amylin and GLP-1 analogues with half-lives on the week timescale, this approach has the potential to provide stable peptide concentrations over several months. This process-level strategy decouples pharmacodynamic optimisation from pharmacokinetic control, providing a generalisable framework to create long-acting, well-tolerated formulations for existing and future peptide therapeutics.

## Methods

### Samples

Pramlintide acetate was purchased from Bachem as lyophilised powder. For biophysical characterisation, peptide stock solutions were prepared immediately prior to use by dissolving the lyophilised peptide in water, filtering through a 0.22 *µ*m syringe filter. The soluble peptide concentration was measured via absorbance at 280 nm (*ε*_280_=1615 M^−1^cm^−1^) and diluted to the final sample concentrations with the relevant buffer stock and water. Buffer solutions were filtered prior to use.

### Thioflavin T fluorescence

Self-assembly was monitored as a function of time using the fluorophore Thioflavin T (ThT), which interacts with *β*-sheet rich fibrils, causing a red-shift in the fluorescence emission spectrum. Fluorescence intensity time courses were recorded on a BMG FluoStar Omega plate reader with a heating function using a 440 nm excitation/ 480 nm emission filter set with 10 nm bandwidths. Time points were acquired at five-minute intervals at 37^◦^C unless otherwise indicated. Measurements were made in black Corning non-binding surface 96-well plates, using sample volumes of 90 *µ*l for forward kinetics in half-area plates (catalogue number 3881) or 200 *µ*l for dissociation kinetics in full area plates (catalogue number 3651). Each sample condition was loaded in triplicates.

The following buffer systems were tested: sodium citrate pH 3; sodium citrate pH 4; sodium acetate pH 4; sodium acetate pH 5, 2-(N-morpholino)ethanesulfonic acid (MES) pH 6; Hepes-NaOH pH 7, Hepes-NaOH pH 7.4. Unless otherwise specified, sample solutions were prepared in 30 mM buffer, 150 mM NaCl, 20 *µ*M ThT, 0.01 % v/v Tween20.

ThT fluorescence curves were fitted using the AmyloFit platform, see Meisl *et al.* Nature Protocols, 2016^37^ for details of the data fitting protocol. We used a model including primary nucleation, elongation and secondary nucleation, with a fixed primary nucleus size *n_c_* = 2. For each buffer condition *n*_2_, *k*_+_*k_n_*, *k*_+_*k*_2_ were fitted as global parameters. Three fluorescence curves were analysed for each sample condition and concentration across ≥6 peptide concentrations (except 3.3 mg/ml peptide in acetate pH 4 and 4.1 mg/ml in citrate pH 4 where n = 2).

### Atomic force microscopy

AFM images of pramlintide were collected using a Bruker Multimode instrument in tapping mode (citrate pH 4, acetate pH 5, MES pH 6, Hepes pH 7, acetate pH 4 and Hepes pH 7.4 cross-dilution matrix) or a Park Instruments NX10 in non-contact mode (citrate pH 3, acetate pH 4, Hepes pH 7.4, 4^◦^C storage time course). Samples were diluted to 0.2-0.5 mg/ml, deposited on freshly cleaved mica and air dried. We used the values for *h* to assess the fibril cross-section, as these are more accurate than the fibril width in the images. Images were analysed in Gwyddion and using the Topostats workflow.^50^

### Pharmacokinetics

Pramlintide self-assembly reactions were carried out at 37^◦^C in sterile solutions of 30 mM sodium acetate pH 4 or 30 mM Hepes-NaOH at pH 7.4 with 150 mM NaCl. Fibrils were stored at 4^◦^C until use. To confirm that the structures would be stable during storage and shipping at 4^◦^, we refrigerated a set of fibril samples for five weeks, characterising them by AFM at *t* = 0, 1 and 5 weeks, SI Figure 12.

Sprague Dawley rats were supplied by Charles River at 200-250 g and housed 3/cage on standard chow diet (Envigo, 2918) and automatic water. After 1 week of acclimation, on day 0 rats were sorted on body weight (BW) by cage. On day 1 of study, rats received a single dose via subcutaneous injection, according to Table 2. Whole blood (125 *µ*l) was collected at the following timepoints: 0hr, 0.5hr, 1hr, 6hr, 1d, 2d, 4d, 7d, 10d, 14d, 22d, and 28d and spun for serum. For the rats receiving IV peptide, blood collections were alternated between individuals 1-3 and 4-6 in each group. Group C only had four rats. At termination, the injection site depot was located where possible, collected and weighed.

### Chromatography

Following aggregation/dissociation, soluble pramlintide was quantified using a Waters Acquity UPLC system. 50% v/v acetonitrile was added to the samples, and these were stored at 10^◦^C during the run sequence. Sample injections of 5 – 10 *µ*l were made on a BEH C18 1.7 *µ*m 2.1 × 100 mm column at 40^◦^C and eluted using an acetonitrile-water gradient (70% to 20% v/v water over four minutes, 0.4 ml/min flow rate, total method duration 6.3 minutes) with a constant trifluoroacetic acid concentration of 0.05% v/v. Peptide backbone absorbance was detected at 220 nm. Peptide concentration was determined based on the peak area relative to a reference curve (linear fit) for standard solutions from 0.001 to 0.5 mg/ml.

For all serum samples, a volume of 50 *µ*l was precipitated with 300 *µ*l cold acetonitrile:methanol (1:1) with 0.2% v/v formic acid. After a brief vortex mix followed by centrifugation (3220 x g, 20 min, at 4^◦^C), 220 *µ*l of the supernatant was evaporated with nitrogen gas to dryness. Samples were reconstituted in 100 *µ*l 25% v/v acetonitrile, 25% v/v methanol with 0.2% v/v formic acid (aq) and vortexed for 15 min prior to injection on the LC-MSMS system.

Pramlintide was analyzed on Waters Xevo TQ-Absolute triple quadrupole mass spectrometer coupled to a Waters Acquity UPLC system, and the chromatografic separation was performed using ACQUITY Peptide BEH C18, 300A, 1.8 *µ*m, 2.1×50 mm column at a temperature of 50^◦^C. Pramlintide was quantified by using mrm transition m/z: 988.4 *>* 968.2.

## Author contributions

TWH, SG, DW, DL, HB, MYA, AB, DH, TPJK, ALGS designed the project. TWH, CR, MP, EB, SW carried out experiments. TWH, JW, PG performed data analysis. All authors contributed to the manuscript.

