## Supplementary Information for "Reversible peptide self-assembly enables sustained drug delivery with tuneable pharmacokinetics"

**Kinetic analysis of the forward self-assembly reaction**

We used the online platform Amylofit to analyse the kinetics of pramlintide self-assembly.[1] To assess the relative importance of primary nucleation versus secondary nucleation driven increases in the fibril population, we compare the two unitless parameters $\lambda = \sqrt{2k_+k_nm^{n_c}}$ and $\kappa = \sqrt{2k_+k_2m^{n_2+1}}$, SI Figure 2**a-b**. This analysis shows that the rate of fibril proliferation through secondary processes increases above pH 5 and drives the observed acceleration in the self-assembly reaction, Figure 2**a-b**.

We measure a relatively low $k_+$ for citrate pH 3, SI Figure 2**d** and SI Figure 4**a**, which is reflected in correspondingly high values for $k_n$ and $k_2$. Due to the shallow slope of the seeded data for citrate pH 3, the rate constants for this buffer system are not as well constrained as those for the remaining solution conditions. In terms of morphology, the fibrils in citrate at pH 3, were shorter and wider than for the other buffer systems investigated here, Figure 2**c-d**. The low observed $k_+$ may therefore be due to steric inhibition. Below, we focus on acetate at pH 4 and Hepes pH 7.4.

### Microfluidic analysis of solution properties

The rate of secondary nucleation $k_2$ is higher in the multivalent citrate buffer than in acetate, Table 1. $k_2$ then increases monotonically for acetate pH 4 through Hepes pH 7.4. The citrate buffer system is a mixture of mono- and divalent ions, and the higher ion valency may provide increased charge screening between peptides. However, the overall ionic strengths of the sample solutions are similar due to the presence of 150 mM NaCl. To investigate whether the buffer systems affected the size and charge of pramlintide in solution, we used microfluidic diffusional sizing and free-flow electrophoresis techniques as reported previously, SI Figure 3.[2–4] The peptide was imaged via intrinsic fluorescence from the tyrosine residue to avoid perturbing the structure with a fluorophore label. We found that at low buffer concentrations (3 mM, no salt), the hydrodynamic radius, $R_H$, and electrophoretic mobility, $\mu_e$, of pramlintide in citrate were larger than would be predicted for even a fully unfolded sequence, indicating that oligomers are formed, whereas the peptide was compacted in the other buffer systems. As we increased the buffer and NaCl concentration to 30 mM and 150 mM respectively, the peptide $R_H$ assumed values of 1.45±0.09 nm to 1.86±0.5 nm, consistent with a partially folded peptide that contains a disulphide bridge.[5]

### Numerical simulation of peptide release from self-assembled depot

We model the serum pramlintide concentration following a single subcutaneous depot injection by combining the $k_+$, $k_-$ and soluble peptide concentrations determined in our *in vitro* analysis,

Table 1. We modelled $M$ as a function of time,[6]

$$M(t) = (m_t - \frac{k_-}{k_+})(1 - e^{-n_{end}k_+t}) + M_0\, e^{-n_{end}k_+t}. \tag{1}$$

In a closed system, the change in $M$ against $t$ is:[6]

$$\frac{dM(t)}{dt} = n_{end} \cdot [k_+(m_t - M(t)) - k_-]. \tag{2}$$

We combined the fit parameters for *in vitro* peptide association and dissociation from fibril structures ($k_+$, $k_-$, endpoint concentrations), SI Figure 2**c**, with the fit parameters for a two-compartment model,[7] and a literature value for the absorption rate from the subcutaneous space to the central compartment, Figure 4**a**.[8] To understand the predictive power of our *in vitro* analysis for subcutaneous depots, we performed numerical simulations for a 300 g rat using the following set of ordinary differential equations to describe peptide movement between states:

$$\frac{dM(t)}{dt} = -k_a\, m_{sc} - \frac{2\, M}{l} \cdot (k_+\, m_{sc} - k_-) \tag{3}$$

$$\frac{dm_{sc}(t)}{dt} = \frac{2\, M}{l} \cdot (k_+\, m_{sc} - k_-) - D_e\, k_e\, M \tag{4}$$

$$\frac{dm_1(t)}{dt} = -k_{12}\, m_1 + k_{21}\, m_2 \cdot \frac{V_2}{V_1} + k_a\, m_{sc} \cdot \frac{V_d}{V_1} - k_e\, m_1 \tag{5}$$

$$\frac{dm_2(t)}{dt} = k_{12}\, m_1 \cdot \frac{V_1}{V_2} - k_{21}\, m_2 \tag{6}$$

$$\frac{dm_e(t)}{dt} = k_e\, m_1 \cdot \frac{V_1}{V_d} + D_e\, k_e\, M \tag{7}$$

Numerical simulations of $m_1$ ($m_{obs}$) were compared with serum pramlintide concentrations following depot administration Figure 4**b-c**. For both the pH 4 and pH 7.4 formulations, we used the $k_+$, $k_-$ values determined for dissociation and growth in pH 7.4 buffer, Figure 3 and SI Figure 4, as these are the closest match to the physiological environment. When we compare simulations using rates for pH 4→4 with pH 4→7.4, SI Figure 8, we find that indeed rates for pH 4→7.4 describe the observed pharmacokinetic profile best.

The value of $D_e$ in this study was constrained by considering the number of rats with detectable depot present at $t$ = 28 days, SI Figure 9. In addition, we considered the predicted dose dependence, SI Figure 10, and the half-time of the $\beta$-phase (HTB), once $m_1$ and $m_2$ have equilibrated, SI Figure 11.[7,9]

$$SUM = k_e + k_{12} + k_{21} \tag{8}$$

$$ROOT = \sqrt{SUM^2 - 4 \cdot K_{21} \cdot k_e} \tag{9}$$

$$\alpha = 0.5 \cdot (SUM + ROOT) \tag{10}$$

$$\beta = 0.5 \cdot (SUM - ROOT) \tag{11}$$

$$HTA = \frac{ln(2)}{\alpha} \tag{12}$$

$$HTB = \frac{ln(2)}{\beta} \tag{13}$$

We adjusted HTB by changing $k_e$ to reflect the increased stability of long-acting peptides.[10] Reducing $k_e$ also decreases the predicted depot clearance, which extends the predicted pharmacokinetics, SI Figure 11. Further testing in *in vivo* systems would be needed to determine the correlation between peptide elimination from the depot in the subcutaneous space and in the central volume.


1. Meisl, G. *et al.* Molecular mechanisms of protein aggregation from global fitting of kinetic models. *Nature Protocols* **11**, 252–272 (2016).

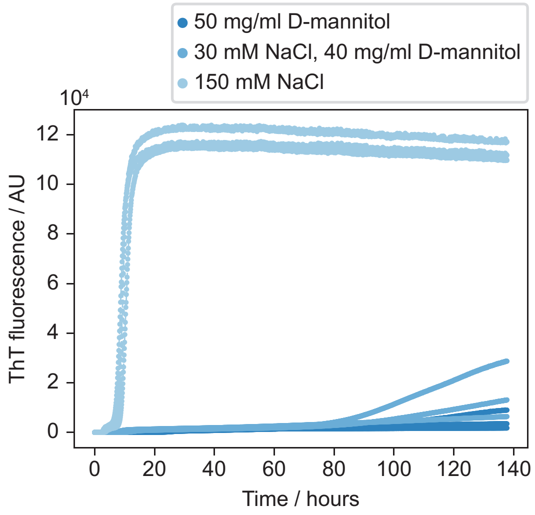


Figure 1: Pramlintide self-assembly accelerates with increasing ionic strength. Pramlintide self-assembly as a function of whether salt (NaCl) or sugar (D-mannitol) is used for isotonicity. 16 mg/ml pramlintide in 30 mM acetate pH 4. No Tween20 was added for this initial screen.


2. Herling, T. W. *et al.* Integration and characterization of solid wall electrodes in microfluidic devices fabricated in a single photolithography step. *Applied Physics Letters* **102**, 184102 (2013).

3. Arosio, P. *et al.* Microfluidic diffusion analysis of the sizes and interactions of proteins under native solution conditions. *ACS NAno* **10**, 333–341 (2016).

4. Herling, T. W. *et al.* Nonspecificity fingerprints for clinical-stage antibodies in solution. *Proceedings of the National Academy of Sciences* **120**, 1–8 (2023).

5. Danielsson, J., Jarvet, J., Damberg, P. & Gräslund, A. Translational diffusion measured by PFG-NMR on full length and fragments of the Alzheimer a$\beta$(1-40) peptide. determination

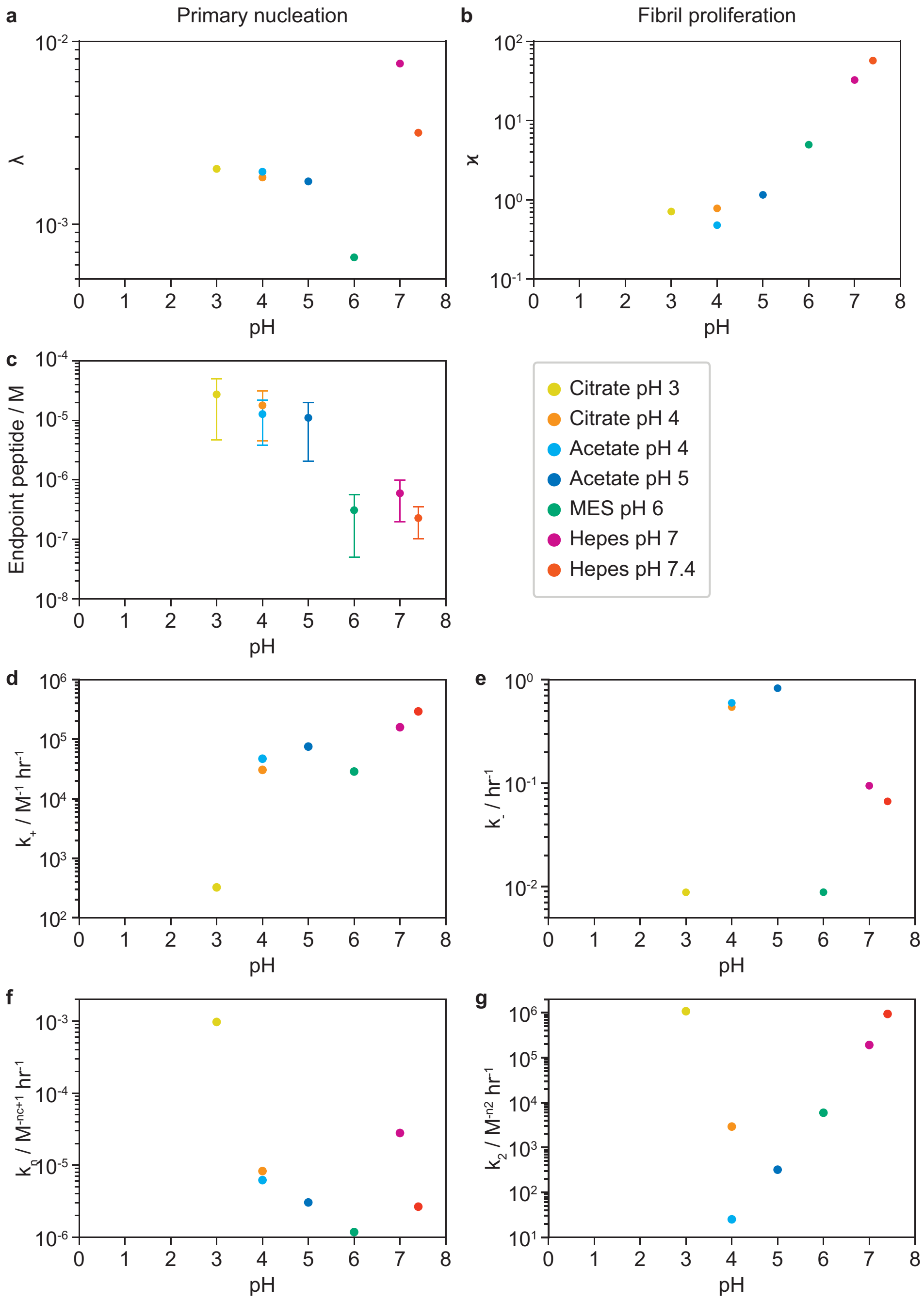


Figure 2: Secondary nucleation and elongation accelerate pramlintide self-assembly with increasing pH. Analysis of $\lambda$ (**a**) and $\kappa$ (**b**) for $m = 10$ mg/ml (2.53 mM) reveal the balance between primary and secondary processes. **c** The endpoint concentration of soluble pramlintide at reaction endpoints (Figure 2) determined by UPLC, error bars show the standard deviation. **d** Elongation rate, $k_+$, from fits to seeded data and endpoint concentration. **e** Dissociation rate $k_-$ based on endpoint peptide and $k_+$. $k_n$ (**f**) and $k_2$ (**g**) as a function of pH from fit to forward kinetics and seeded reactions.

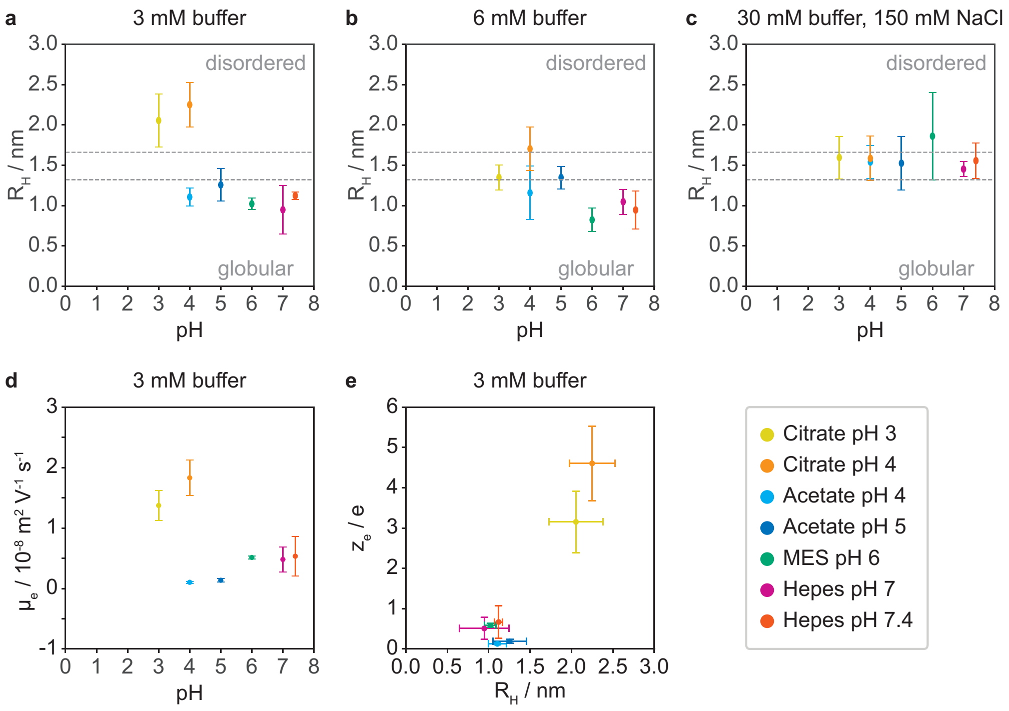


Figure 3: Size and charge of pramlintide are buffer dependent at low ionic strength. **a-c** $R_H$ from microfluidic diffusional sizing of 10 mg/ml pramlintide at increasing buffer and salt concentrations. The dashed lines represent the expected $R_H$ for a disordered and a globular protein with the molecular weight of pramlintide.[5] The size of the peptide is highly buffer dependent at low ionic strength, but converges to the expected value for a partially unfolded protein at physiological ionic strength levels. Unlabelled peptide was imaged via intrinsic fluorescence. **d** $mu_e$ for 10 mg/ml pramlintide in 3 mM buffer. The low conductivity of the acetate buffers at pH 4 and 5 lead to low apparent $\mu_e$ values. **e** Effective charge against $R_H$ in 3 mM buffer. Pramlintide in citrate is both larger and carries a higher overall charge than in the other buffer systems.

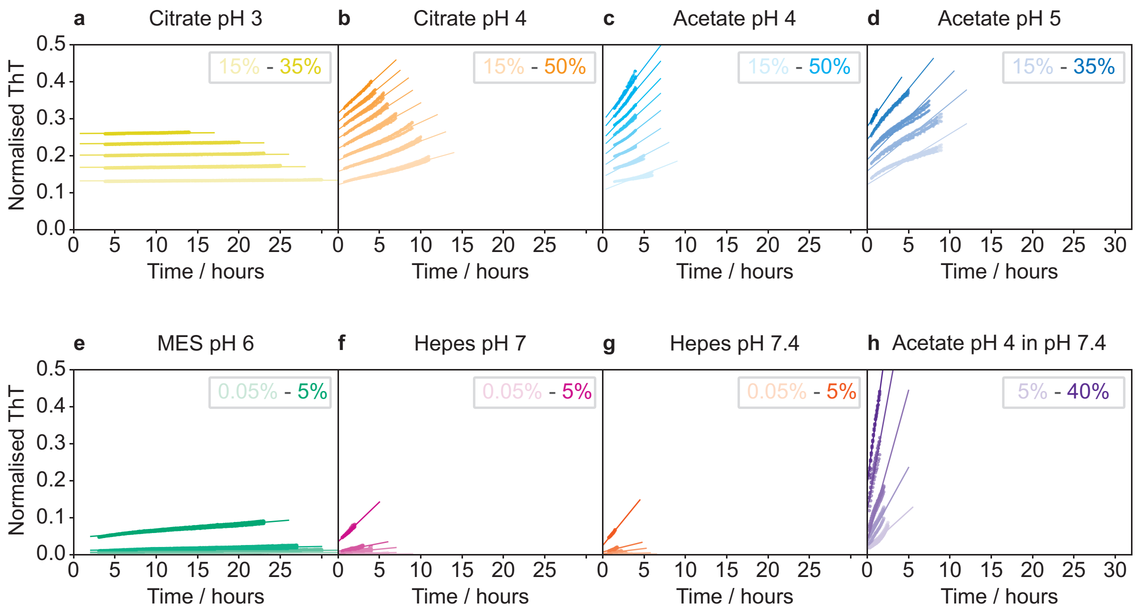


Figure 4: Pramlintide self-assembly $k_+$ increases with pH. 1 mg/ml solutions of pramlintide (253 $\mu$M) were seeded with preformed fibrils and the elongation rate ($k_+$) determined from global fits to the linear sections of ThT fluorescence curves for each buffer. Here, seed elongation is the primary process, before secondary nucleation dominates the fluorescence intensity increase.[11,12] The seed concentration is expressed as a percentage of the monomer e.g. 5% for 1 mg/ml monomer plus 0.05 mg/ml monomer equivalent in seed fibril. For samples in panels **a-d** the seed concentration was increased in 5% increments. In panels **e-g**, 0.05, 0.1, 0.5, 1, and 5% seed were added. **h** Cross-seeding of a pH 7.4 solution with fibrils formed in acetate pH 4. 5, 10, 20, 30, 40% seed were added.

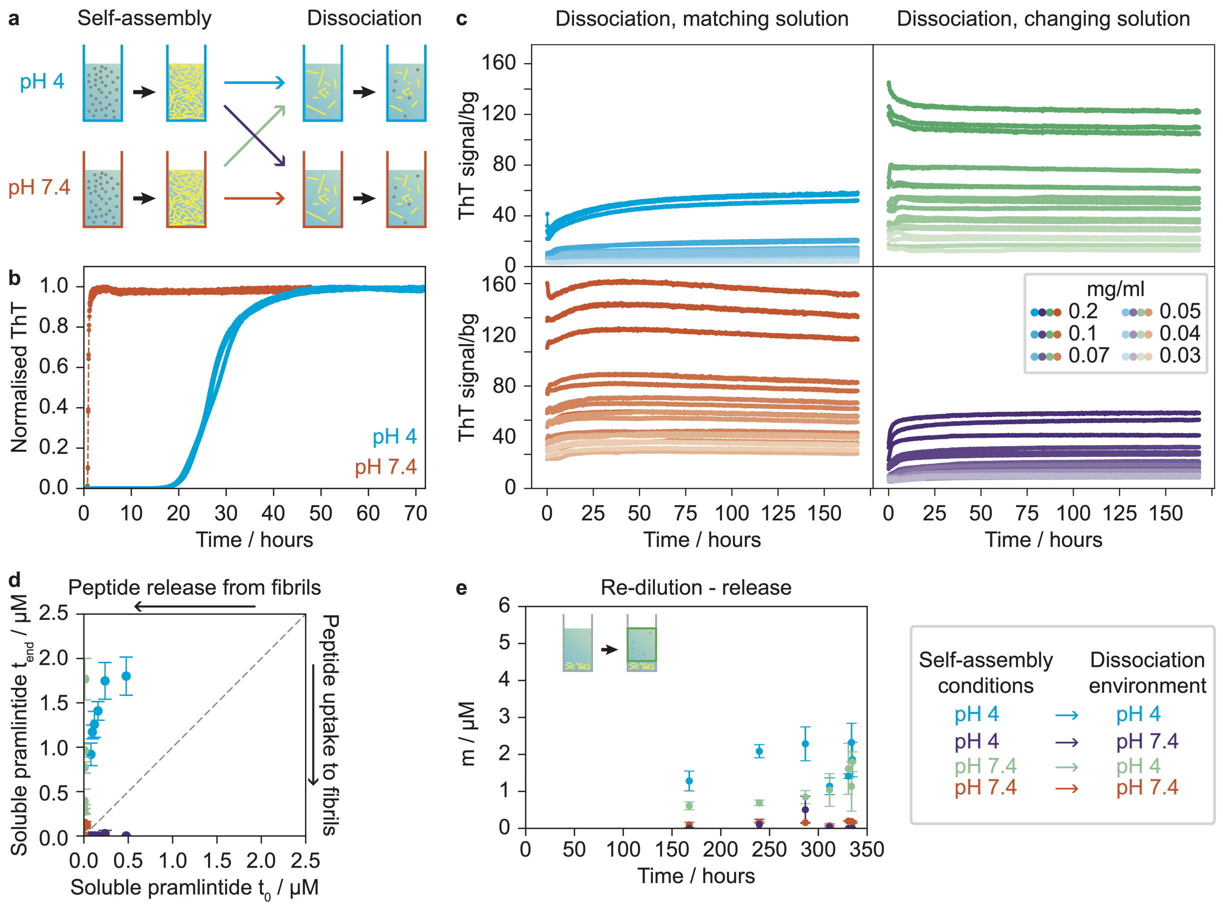


Figure 5: Tuneable dissociation kinetics for pramlintide. **a** Schematic of experiment workflow. **b** 10 mg/ml pramlintide is self-assembled in acetate pH 4 or Hepes pH 7.4, both with 150 mM NaCl, 20 $\mu$M ThT, 0.01% v/v Tween20. **c** The two fibril types are then diluted in either pH 4 or pH 7.4 buffer to create four sample sets, two with the dilution buffer is different from the self-assembly buffer, and two where the solution conditions match and monitored by ThT fluorescence for seven days. The legend in the bottom right corner of the figure applies to all panels. **d** Pramlintide is released or taken up to fibrils following dilution, depending on the combination of self-assembly and dissociation environments. The diagonal line corresponds to no uptake/release of peptide, t = 7 days after dilution. **e** Soluble pramlintide as a function of time following removal and replacement of the soluble fraction in Figure 3**d**.

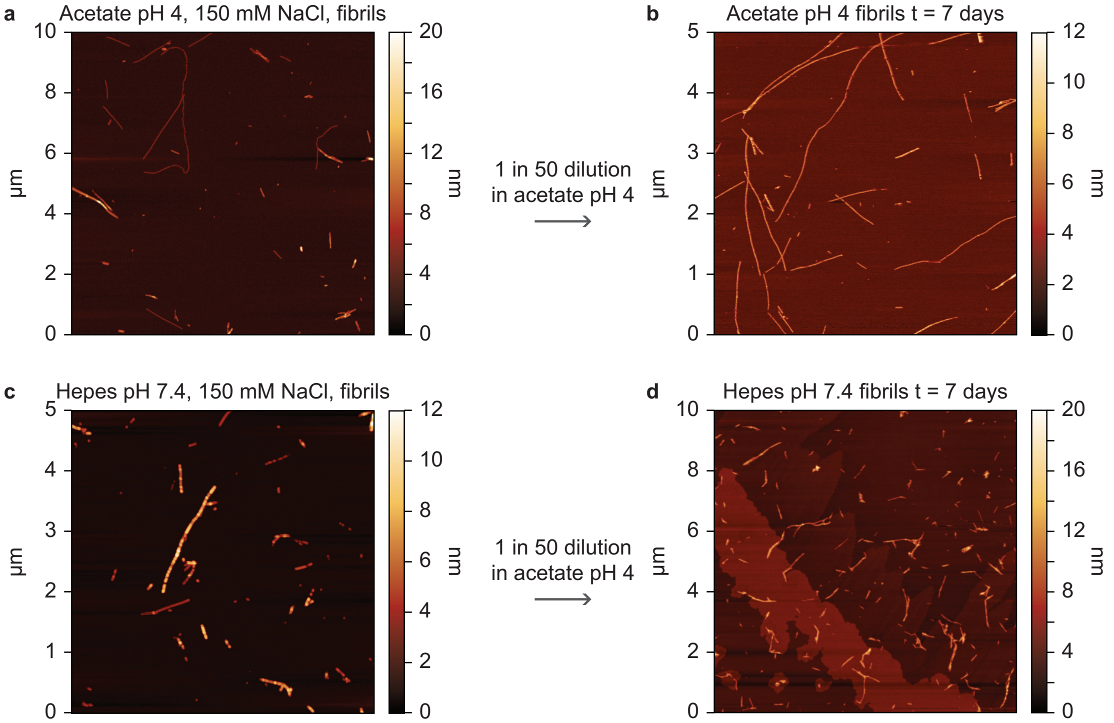


Figure 6: Fibrils retain distinct morphologies after dilution in different buffer systems. **a** Fibrils made in 30 mM acetate pH 4, 150 mM NaCl prior to dilution (t = 0 days). **b** Fibrils at t = 7 days after dilution and storage in 30 mM acetate pH 4, 0.02% w/v $NaN_3$, 20 $\mu$M ThT. Note the difference in image area between panels. **c** Fibrils made in 30 mM Hepes pH 7.4, 150 mM NaCl prior to dilution (t = 0 days). **d** Fibrils at t = 7 days after dilution and storage in 30 mM acetate pH 4, 0.02% w/v $NaN_3$, 20 $\mu$M ThT.

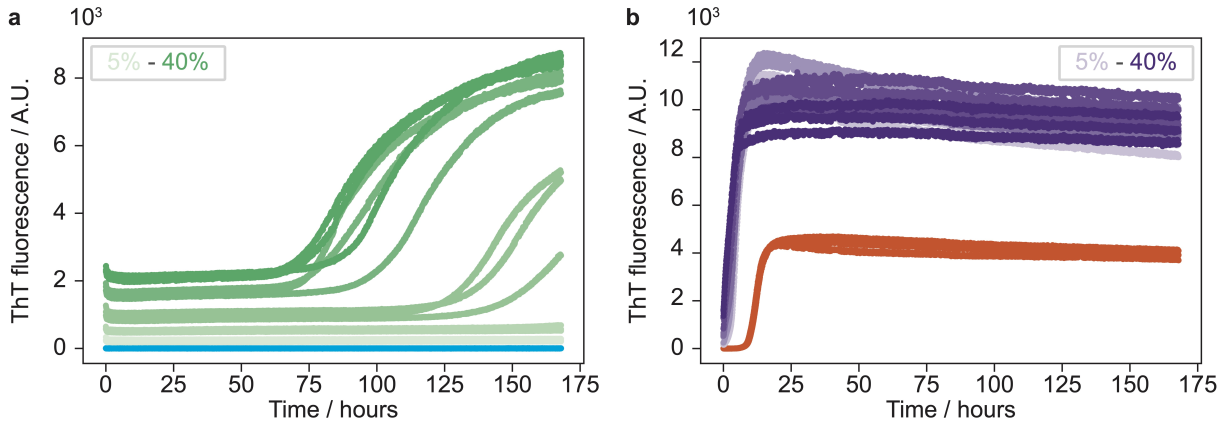


Figure 7: Cross-seeding efficiency depends on the solution conditions. **a** 1 mg/ml solutions of pramlintide (253 $\mu$M) in acetate pH 4 were seeded with preformed full length fibrils prepared in the Hepes pH 7.4 buffer system. Blue is 0% seed, for the green data 5, 10, 20, 30, 40% seed were added. **b** Cross-seeding of a pH 7.4 solution with fibrils formed in acetate pH 4, vermillion data is unseeded pramlintide. 5, 10, 20, 30, 40% seed were added (purple data points).

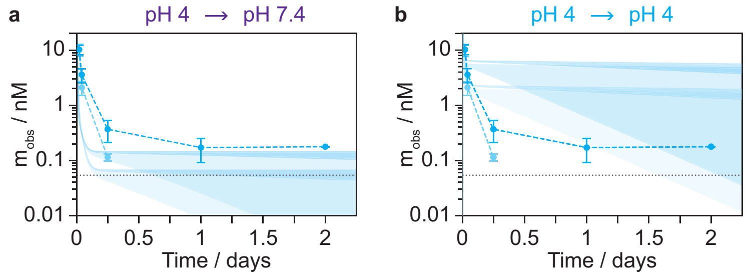


Figure 8: Rate constants for transfer to pH 7.4 buffer agree with pharmacokinetic data. **a** Serum pramlintide with $k_+$, $k_-$ from pH 4→7.4 buffer and $D_e$ of 0 – 0.001. **b** Serum pramlintide with $k_+$, $k_-$ from pH 4→4 buffer and $D_e$ of 0 to $10^{-4}$ (dark) and $10^{-3}$ (light).

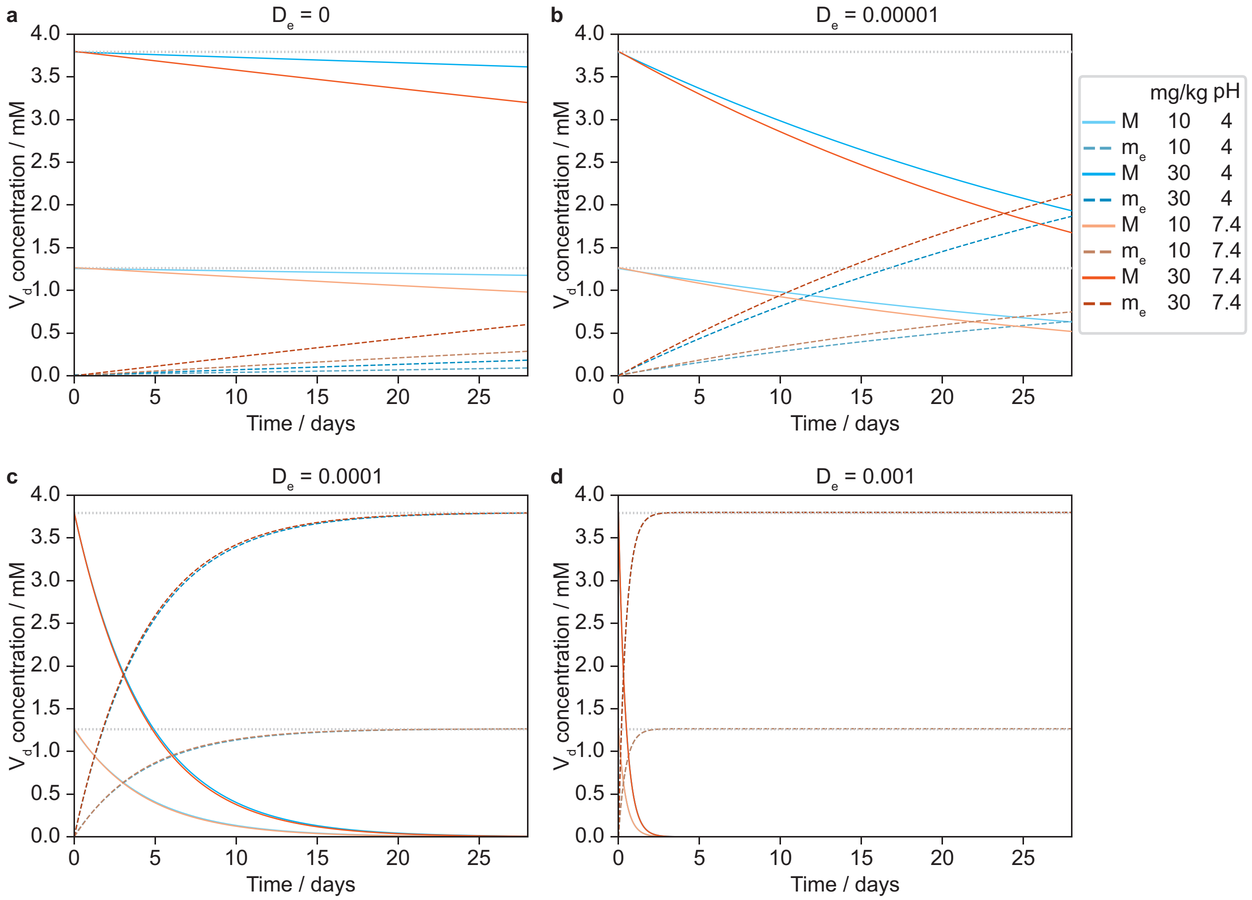


Figure 9: The effect of $D_e$ on self-assembled depot depletion. **a** Depot concentration as a function of time without elimination via the subcutaneous space ($D_e$ = 0) and eliminated peptide for $V = V_d$. **b** $M$ and $m_e$ as a function of time for $D_e = 10^{-5}$. **c** $M$ and $m_e$ as a function of time for $D_e = 10^{-4}$. **d** $M$ and $m_e$ as a function of time for $D_e = 10^{-3}$.

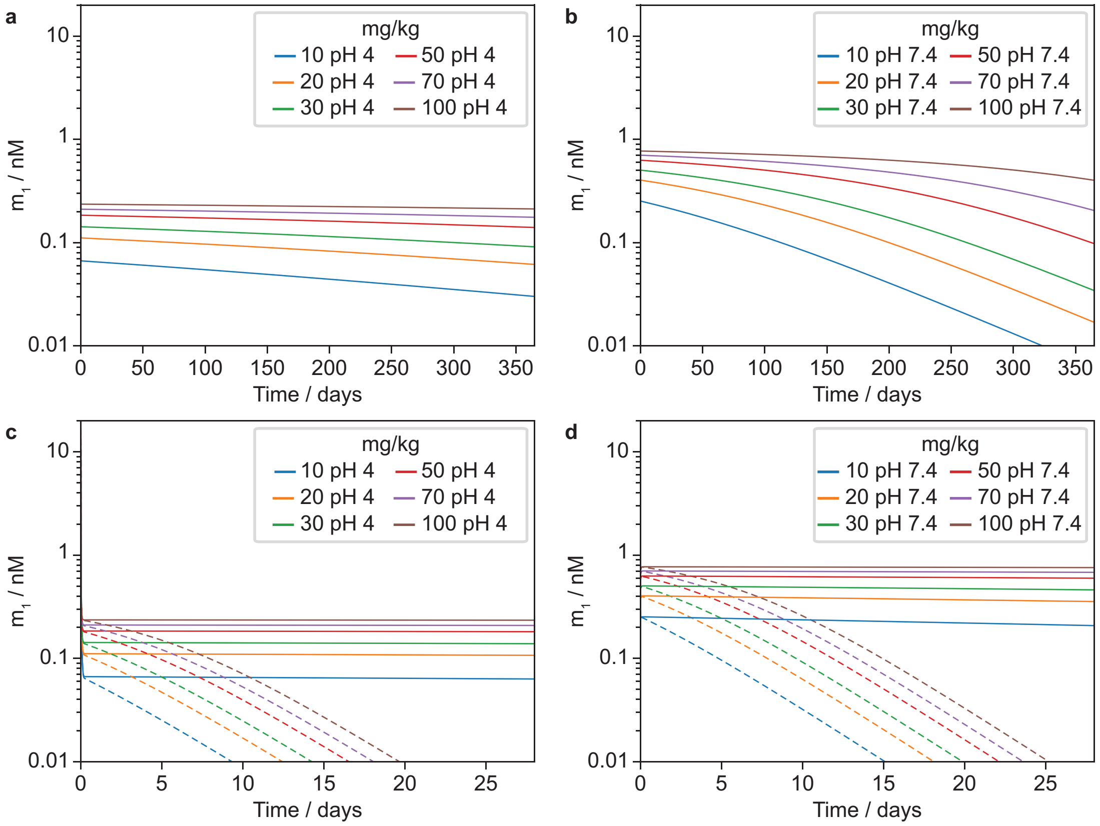


Figure 10: Simulation of dose dependence of peptide exposure aids formulation design. **a** Predicted $m_1$ for pH 4 formulations as a function of dose for $D_e$ = 0 over long timescales. **b** Predicted $m_1$ for pH 7.4 formulations as a function of dose for $D_e$ = 0 over long timescales. During the steady state phase, $m_1$ is higher as a function of dose for pH 7.4. Both $k_-$ and $n_{end}$ are lower for pH 4 formulations, leading to a longer steady state for pH 4 depots. **c** Predicted $m_1$ for pH 4 formulations in rat as a function of dose for $D_e$ = 0 (solid lines) and $D_e$ = 0.0001 (dashed lines). **d** Predicted $m_1$ for pH 7.4 formulations as a function of dose for $D_e$ = 0 (solid lines) and $D_e$ = 0.0001 (dashed lines).

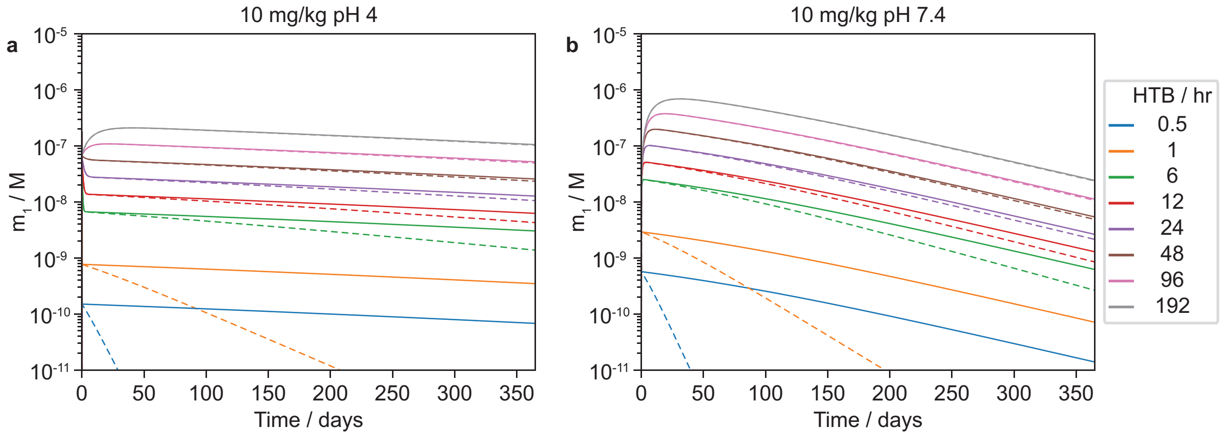


Figure 11: Simulations of serum peptide concentrations indicate that long-acting peptides enable monthly dosage. **a** Predicted $m_1$ for pH 4 formulations as a function of the half-life of the $\beta$ phase of a two-compartment model. The peptide half-life is regulated by changing $k_e$. In this model, the rate of depot clearance in the subcutaneous space is proportional to that for circulating peptide in $V_1$. Here, $m_1$ is simulated for $D_e$ = 0 (solid lines) and $D_e$ = 0.0001 (dashed lines). **b** Predicted $m_1$ for pH 7.4 formulations as a function of the half-life of the $\beta$ phase of a two-compartment model.

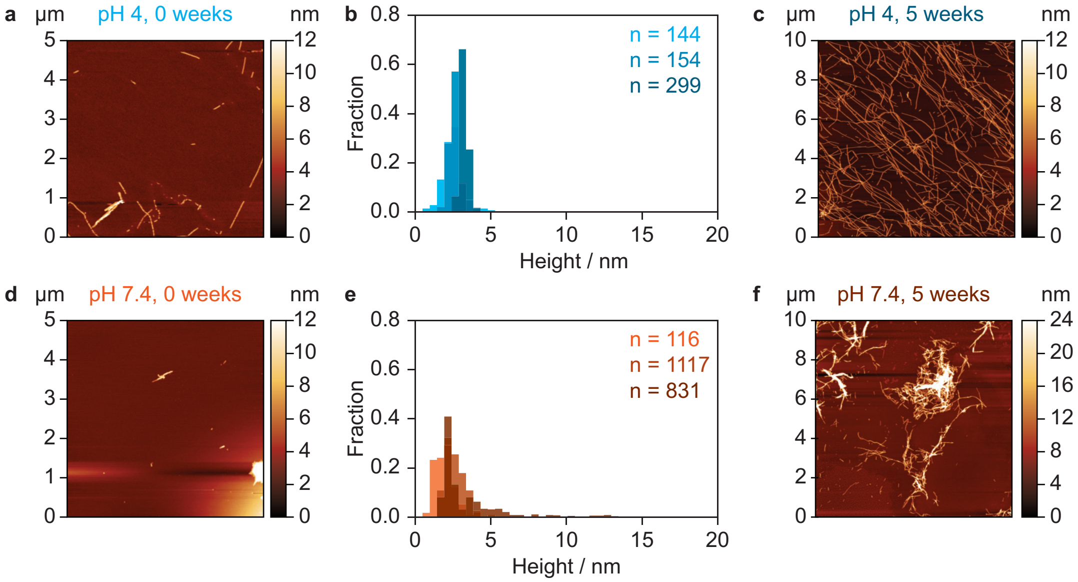


Figure 12: Fibril stability and development during storage at 4°C. **a** Self-assembled pramlintide prepared in acetate pH 4 at 37°C. **b** Time evolution of fibril height when stored at 4°C for sample in panel **a**. The fibril thickness (height) increases slightly over time, as the structures mature to form networks of long fibrils. **c** Acetate pH 4 samples after five weeks of storage. **d** Self-assembled pramlintide prepared in Hepes pH 7.4 at 37°C. **e** Time evolution of height distribution during storage at 4°C for sample in panel **d**. During storage, the pH 7.4 fibrils mature towards a more uniform height distribution within the initial range. **f** AFM images after five weeks at 4°C shows that the pH 7.4 fibrils form clusters, but do not mature to the multi-$\mu$m long structures seen at pH 4.

of hydrodynamic radii of random coil peptides of varying length. *Magnetic Resonance in Chemistry* **40**, S89–S97 (2002).

6. Knowles, T. P. J. *et al.* An analytical solution to the kinetics of breakable filament assembly. *Science* **326**, 1533–1537 (2009).

7. Nordell, P., Jansson-Löfmark, R. & Gennemark, P. Systemic pharmacokinetic principles of therapeutic peptides. *Clinical Pharmacokinetics* (2026).
